# Integrated analyses of early responses to radiation in glioblastoma identify new alterations in RNA processing and candidate target genes to improve treatment outcomes

**DOI:** 10.1101/863852

**Authors:** Saket Choudhary, Suzanne C. Burns, Hoda Mirsafian, Wenzheng Li, Dat T. Vo, Mei Qiao, Andrew D. Smith, Luiz O. Penalva

## Abstract

**Background:** High-dose radiation is the main component of glioblastoma therapy. Unfortunately, radio-resistance is a common problem and a major contributor to tumor relapse. Understanding the molecular mechanisms driving response to radiation is critical for identifying regulatory routes that could be targeted to improve treatment response.

**Methods:** We conducted an integrated analysis in the U251 and U343 glioblastoma cell lines to map early alterations in the expression of genes at three levels: transcription, splicing, and translation in response to ionizing radiation.

**Results:** Changes at the transcriptional level were the most prevalent response. Downregulated genes are strongly associated with cell cycle and DNA replication and linked to a coordinated module of expression. Alterations in this group are likely driven by decreased expression of the transcription factor FOXM1 and members of the E2F family. Genes involved in RNA regulatory mechanisms were affected at the mRNA, splicing, and translation levels, highlighting their importance in radiation-response. We identified a number of oncogenic factors, with an increased expression upon radiation exposure, including BCL6, RRM2B, IDO1, FTH1, APIP, and LRIG2 and lncRNAs NEAT1 and FTX. Several of these targets have been previously implicated in radio-resistance. Therefore, antagonizing their effects post-radiation could increase therapeutic efficacy.

**Conclusions:** Our integrated analysis provides a comprehensive view of early response to radiation in glioblastoma. We identify new biological processes involved in altered expression of various oncogenic factors and suggest new target options to increase radiation sensitivity and prevent relapse.

## Background

Glioblastoma is the most common intracranial malignant brain tumor with an aggressive clinical course. Standard of care entails maximally safe resection followed by radiotherapy with concomitant and adjuvant temozolomide. Nonetheless, the median overall survival remains approximately 16 months [1, 2], and the recent addition of tumor-treating fields to the standard of care has only increased median overall survival to 20.5 months [1]. Recurrence occurs in part because glioblastoma uses sophisticated cellular mechanisms to repair DNA damage from double-stranded breaks caused by ionizing radiation, specifically homologous recombination and non-homologous end-joining. Thus, the repair machinery confers a mechanism for resistance to radiation therapy. Ionizing radiation can also cause base damage and single-strand breaks, which are repaired by base excision and single-strand break repair mechanisms, respectively [3]. A comprehensive analysis of molecular mechanisms driving resistance to chemotherapy and radiation is required to surpass major barriers and advance treatments for glioblastoma.

The Cancer Genome Atlas (TCGA) was instrumental in improving the classification and identification of tumor drivers [4], but its datasets provide limited opportunities to investigate radiation response. Thus, studies using cell and murine models are still the best alternatives to evaluate radiation response at the genomic level. The list of biomarkers associated with radiation resistance in glioblastoma is still relatively small. Among the most relevant are FOXM1 [5, 6], STAT3 [6], L1CAM [7], NOTCH1 [8], RAD51 [9], EZH2 [10], CHK1/ATR [11], COX-2 [12], and XIAP [13]. Dissecting how gene expression is altered by ionizing radiation is critical to identify possible genes and pathways that could increase radio-sensitivity. A few genomic studies [14, 15, 16] have explored this question, but these analyses were restricted to describing changes in transcription.

Gene expression is regulated at multiple levels, and RNA-mediated mechanisms such as splicing and translation are particularly relevant in cancer biology. A growing number of inhibitors against regulators of splicing and translation are being identified [17]. Splicing alterations are a common feature across cancer types and affect all hallmarks of cancer [18]. Numerous splicing regulators display altered expression in glioblastoma (e.g. PTBP1, hnRNPH, and RBM14) and function as oncogenic factors [19]. Importantly, a genome-wide study using patient-derived models revealed that transformation-specific depended on RNA splicing machinery. The SF3b-complex protein PHF5A was required for glioblastoma cells to survive, but not neural stem cells (NSCs). Moreover, genome-wide splicing alterations after PHF5A loss appear only in glioblastoma cells [20]. Translation regulation also plays a critical role in glioblastoma development. Many translation regulators such as elF4E, eEF2, Musashi1, HuR, IGF2BP3, and CPEB1 promote oncogenic activation in glioblastoma, and pathways linked to translation regulation (e.g., mTOR) promote cancer phenotypes [21].

To elucidate expression responses to radiation, we conducted an integrated study in U251 and U343 glioblastoma cell lines covering transcription (mRNAs and lncRNAs), splicing, and translation. We determined that the downregulation of FOXM1 and members of the E2F family are likely the major drivers of observed alterations in cell cycle and DNA replication genes upon radiation exposure. Genes involved in RNA regulatory mechanisms were particularly affected at the transcription, splicing, and translation levels. In addition, we identified several oncogenic factors and genes associated with poor survival in glioblastoma that displayed increased expression upon radiation exposure. Importantly, many have been implicated in radio-resistance, and therefore, their inhibition in combination with radiation could increase therapy efficacy.

## Methods

### Cell culture and radiation treatment

U251 and U343 cells were obtained from the University of Uppsala (Sweden) and maintained in Dulbecco’s Modified Eagle Medium (DMEM, Hyclone) supplemented with 10% fetal bovine serum, 1% Penicillin/Streptomycin at 37°C in 5% CO_2_-humidified incubators and were sub-cultured twice a week. Cells were plated after appropriate dilution, and ionizing radiation treatment was performed on the next day at a dose of 5 Gray (Gy). A cabinet X-ray system (CP-160 Cabinet X-Radiator; Faxitron X-Ray Corp., Tucson, AZ) was used. After exposure to ionizing radiation, cells were cultured for 1 and 24 hours (hrs).

### RNA preparation, RNA-seq and Ribosome Profiling (Ribo-seq)

RNA was purified using a GeneJet RNA kit from Thermo Scientific. The TruSeq Ribo Profile (Mammalian) kit from Illumina was used to prepare material for ribosome profiling (Ribo-seq). RNA-seq and Ribo-seq samples were prepared according to Illumina protocols and sequenced at UTHSCSA Genome Sequencing Facility.

### Overall strategy to identify gene expression alterations upon radiation

To identify the most relevant expression alterations in the early response to radiation, we analyzed samples from U251 and U343 cells collected at 0 (T0), 1 (T1), and 24 (T24) hours post-radiation. To capture the progressive dynamics of expression alterations, we compared T0 to T1 samples and T1 to T24 samples.

Our strategy to identify the most relevant alterations in expression with maximal statistical power was to combine all samples and use a design matrix with cell type defined as a covariate with time points (Figure S1).

### Sequence data pre-processing and mapping

The quality of raw sequences reads from RNA-Seq and Ribo-Seq datasets were assessed using FastQC [22]. Adaptor sequences and low-quality score (phred quality score < 5) bases were trimmed from RNA-Seq and Ribo-Seq datasets with TrimGalore (v0.4.3) [23]. The trimmed reads were then aligned to the human reference genome sequence (Ensembl GRCh38.p7) using STAR aligner (v.2.5.2b) [24] with GENCODE [25] v25 as a guided reference annotation, allowing a mismatch of at most two positions. All the reads mapping to rRNA and tRNA sequences were filtered out before downstream analysis. Most reads in the Ribo-seq samples mapped to the coding domain sequence (CDS). The distribution of fragment lengths for ribosome foot-prints was enriched in the 28-30 nucleotides range, as expected (Figure S2). The ribosome density profiles exhibit high periodicity as within the CDS, as expected since ribosomes traverse three nucleotides at a time (Figure S3). The periodicity analysis was performed using ribotricer [26]. The number of reads assigned to annotated genes included in the reference genome was obtained by htseq-count [27].

### Differential gene expression analysis

For differential expression analysis, we performed counting over exons for the RNA-seq samples. For translational efficiency analyses, counting was restricted to the CDS. A Principal Component Analysis (PCA) was then performed on RNA-Seq and Ribo-Seq data from U251 and U343 cells. Most variation was explained by the cell type along the first principal component, and radiation time-related changes were captured along the second principal component (Supplementary Figure S1B). Differential gene expression analysis was performed by employing the DESeq2 package [28], with read counts from both U251 and U343 cell samples as inputs. We adjusted p-values controlling for the false discovery rate (adjusted p-value) using the Benjamini and Hochberg (BH) procedure [29]. Differentially expressed genes were defined with an adjusted p-value < 0.05

### Weighted gene co-expression network analysis

Weighted Gene Co-expression Network Analysis (WGCNA) [30] uses pairwise correlations on expression values to identify genes significantly co-expressed across samples. We used this approach to identify gene modules with significant co-expression variations as an effect of radiation. The entire set of expressed genes, defined here as those with one or higher transcripts per million higher (TPM), followed by variance stabilization) from U251 and U343 samples were clustered separately using the signed network strategy. We used the *Z*_summary_ [31] statistic as a measure of calculating the degree of module preservation between U251 and U343 cells. *Z*_summary_ is a composite statistic defined as the average of the density and connectivity based statistic. Thus, both density and connectivity are considered for defining the preservation of a module. Modules with *Z*_summary_ > 5 were considered as significantly preserved. The expression profile of all genes in each co-expression module can be summarized as one “eigengene”. We used the eigengene-based connectivity (kME) defined as the correlation of a gene with the corresponding module eigengene to assess the connectivity of genes in a module. The intramodular hub genes were then defined as genes with the highest module membership values (kME >0.9). All analysis was performed using the R package WGCNA. The protein-coding hub genes were then selected for gene ontology enrichment analysis.

### Translational efficiency analysis

We used Riborex [32] to perform differential translational efficiency analysis. The underlying engine selected was DESeq2 [28]. DESeq2 estimates a single dispersion parameter per gene. However, RNA-Seq and Ribo-Seq libraries can have different dispersion parameters owing to different protocols. We estimated the dispersion parameters for RNA-Seq and Ribo-Seq samples separately and found them to be significantly different (mean difference = 0.04, p-value < 2.2*e* – 16). This leads to a skew in translational-efficiency p-value distribution since the estimated null model variance for the Wald test is underestimated. To address this issue, we performed a p-value correction using fdrtool [33] that re-estimates the variance using an empirical bayes approach.

### Alternative splicing analysis

Alternative splicing analysis was performed using rMATS [34]. All reads were trimmed using cutadapt [35] with parameters (`-u −13 -U −13`) to ensure trimmed reads had equal lengths (138 bp). rMATs was run with default parameters in paired end mode (-t paired) and read length set to 138 bp (`-len 138`) using GENCODE GTF (v25) and STAR index for GRCh38.

### Gene ontology (GO) and pathway enrichment analysis

To classify the functions of differentially enriched genes, we performed GO enrichment, and the Reactome pathway [36] analysis using Panther [37]. For both analyses, we considered terms to be significant if BH adjusted p-values weree < 0.05, and fold enrichment is > 2.0. Further, we used REVIGO [38] to reduce redundancy of the enriched GO terms and visualize the semantic clustering of the identified top-scoring terms. We used STRING database (v10) [39] to construct protein-protein interaction networks and determine associations among genes in a given dataset. The interactions are based on experimental evidence procured from high-throughput experiments text-mining, and co-occurrence. Only high-confidence (0.70) nodes were retained.

### Expression correlation analysis

Gene expression correlation analysis was done using Gliovis [40] using glioblastoma samples (RNAseq) from the TCGA. To select correlated genes, we used Pearson correlation, *R* > 0.3, and p-value < 0.05. A list of genes affecting survival in glioblastoma was downloaded from GEPIA [41]. A list of long non-coding RNAs (lncRNAs) implicated in glioma development was obtained from Lnc2Cancer [42]. Drug-gene interactions were identified using the Drug-Gene Interaction Database [43].

## Results

### Changes in global transcriptome profile in response to radiation

We first conducted an integrated analysis to evaluate the early impact of radiation [1 hour (T1) and 24 hours (T24)] on the expression profile of U251 and U343 GBM lines. A relatively small number of genes displayed altered expression at T1. Downregulated genes are mainly involved in transcription regulation and include 18 zinc finger transcription factors displaying high expression correlation in glioblastoma samples from TCGA (Table S1). Upregulated sets contain genes implicated in cell cycle arrest, apoptosis, and stress such as ZFP36, FBXW7, SMAD7, BTG2, and PLK3 (Table S1).

Since many alterations were observed when comparing the T1 vs. T24 time points (Table S1), we opted to focus on genes showing the most marked changes (*log*_2_ fold-change > 1.0 or < −1.0 and adjusted p-value < 0.05) to identify biological processes and pathways most affected at the T24 time point. Top enriched GO terms and pathways among downregulated genes include chromatin remodeling, cell cycle, DNA replication, and repair (Figure 1A). Additionally, we identified several GO terms associated with mRNA metabolism, decay, translation, and ncRNA processing, suggesting active participation of RNA-mediated processes in radio-response (Figure 1B). Network analysis indicated the set of genes in these categories is highly interconnected (Figure 1C and Table S2).

**Figure 1.**
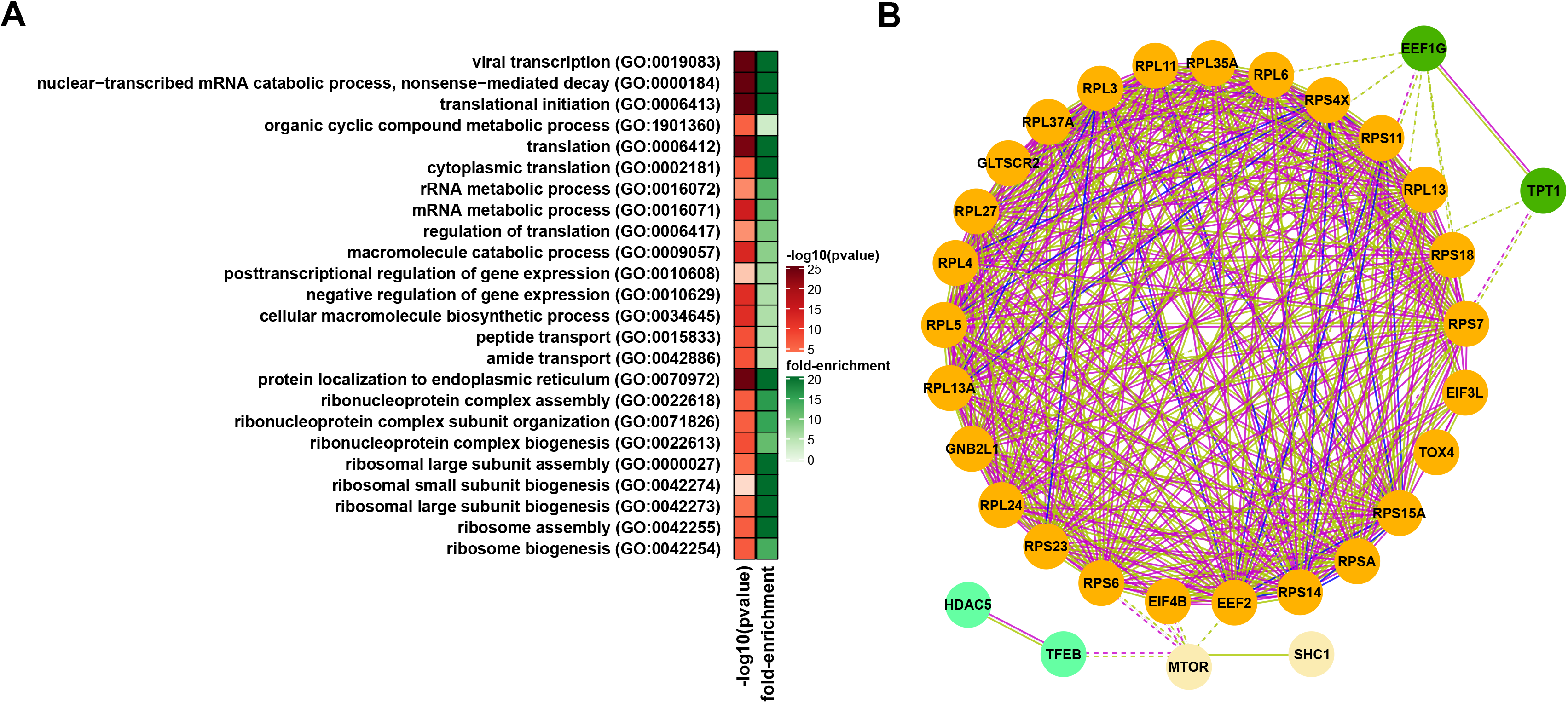
Characteristics of downregulated genes at 24 hours (T24) after radiation exposure in glioblastoma cell lines. A) Enriched gene ontology related to cell cycle, DNA replication, and repair among downregulated genes. B) RNA-related Gene Ontology (GO) terms enriched among downregulated genes summarized using REVIGO [38]. C) Protein-protein interaction network, according to STRING [39] showing downregulated genes associated with RNA-related functions. Gene clusters based on the strength of connection and gene function are identified by color. Lines colors indicate the type of association: light green indicates an association based on literature findings; blue indicates gene co-occurrence; magenta indicates experimental evidence.

To expand the expression analysis, we employed WGCNA [30] to identify gene modules with significant co-expression variation as an effect of radiation. All identified modules, along with the complete list of genes in each module, are shown in Supplementary Figure S4 and Table S3. Seven modules were identified (*Z*_summary_ > 5) as tightly regulated, independent of the cell line (Figure S4E). Among modules with the highest significant correlation (0.8, p-value< 1*e*−7), module 2 contains genes downregulated in T24, with many involved in cell cycle, metabolism mRNA met abolism, processing, splicing, and transport (Table S3), corroborating results described above.

Next, we investigated downregulated genes with the gene set enrichment analysis (GSEA) tool Enrichr [44] and conducted expression correlation analysis with Gliovis [40]. Based on their genomic binding profiles and effect of gene expression, FOXM1 and the E2F family of transcription factors emerged as potential regulators of a large group of cell cycle/DNA replication-related genes in the affected set (Figure 2A, Table S4). In agreement, E2F1, E2F2, E2F8, and FOXM1 displayed a significant decrease upon radiation. FOXM1 and E2F factors have been previously implicated in chromatin remodeling, cell cycle regulation, DNA repair, and radio-resistance [45, 46]. All four factors are highly expressed in glioblastoma with respect to low-grade glioma. Importantly, they display high expression correlation with a large set of down-regulated genes implicated in cell cycle and DNA replication and among themselves in glioblastoma samples in TCGA (Figure 2B-C).

**Figure 2.**
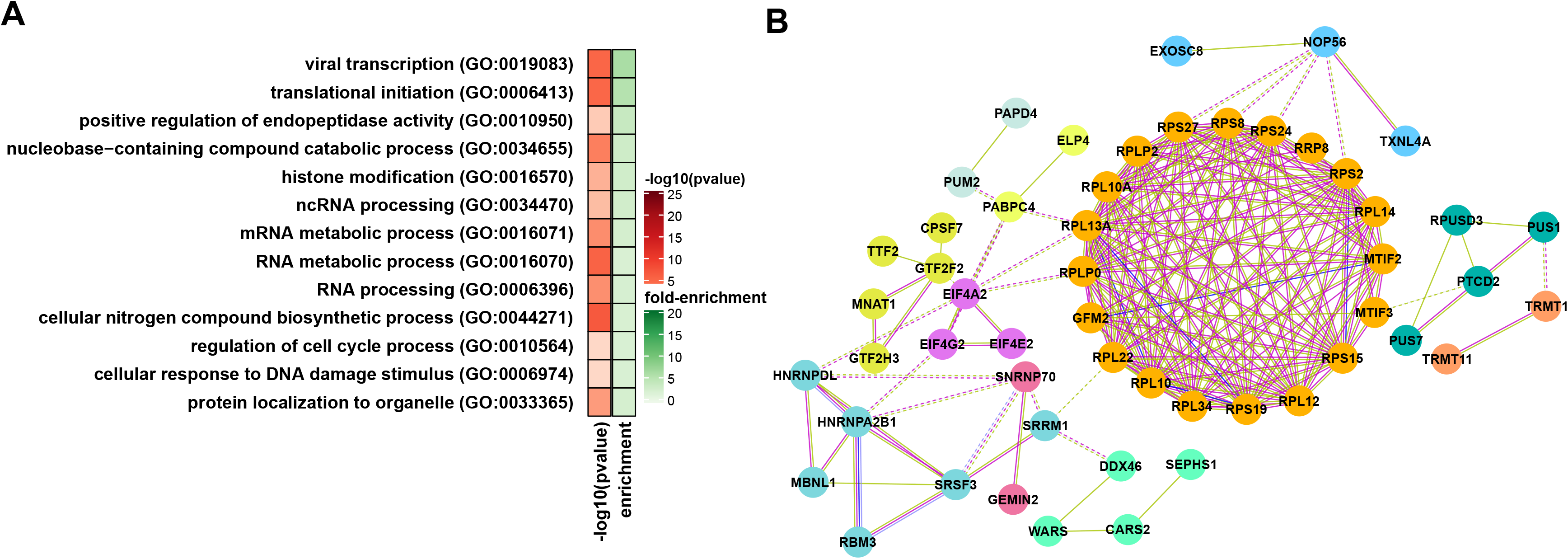
E2Fs and FOXM1 in glioblastoma. A) Correlation of E2F1, E2F2, E2F8, and FOXM1 with target genes involved in cell cycle. B) Expression levels of E2F1, E2F2, E2F8, and FOXM1 in gliomas grades II, III, and IV in TCGA samples. C) E2F1, E2F2, E2F8, and FOXM1 expression correlation in glioblastoma (TCGA samples) using Gliovis [40]. *** p-value < 0.0001.

Upregulated genes at T24 are preferentially associated with the extracellular matrix receptor interaction pathway, extracellular matrix organization, axonogenesis, and response to type I interferon (Figure 3A, and Table S2). With respect to the extracellular matrix, we observed changes in the expression levels of several collagens (types II, IV, V, and XI), glycoproteins of the laminin family (subunits *α, β*, and *γ*), and also integrins (subunits *α*, and *β*) (Figure 3B, and Table S1). Collagen type IV is highly expressed in glioblastoma and implicated in tumor progression [47]. In addition, it has been observed that the activation of two integrins, ITGB3 and ITGB5, contributes to radio-resistance [48].

**Figure 3.**
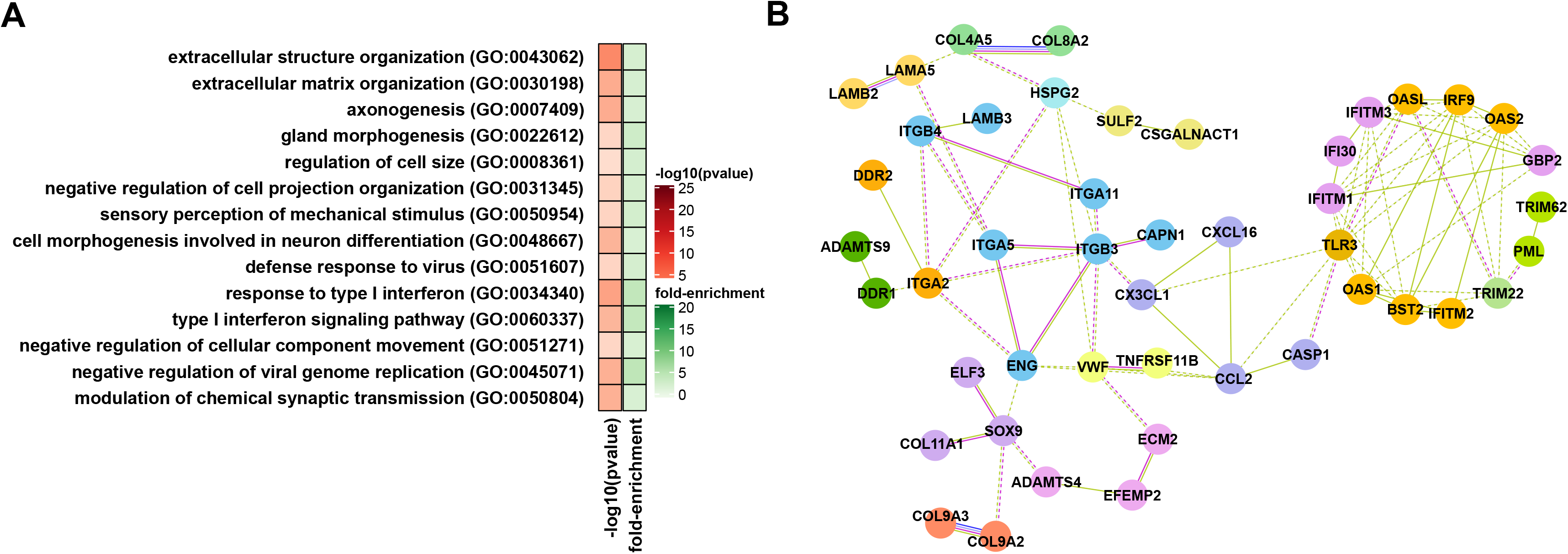
Global view of upregulated genes at T24 post-radiation in glioblastoma cells. A) Gene ontology analysis of upregulated genes B) Protein-protein interaction networks according to STRING [39] showing genes associated with extracellular matrix organization and response to interferon. Gene clusters based on the strength of connection and gene function are identified by color. Lines colors indicate type of association: light green, association based on literature findings; blue indicates gene co-occurrence; magenta indicates experimental evidence.

Radiation treatment also induced the expression of genes involved in neuronal differentiation and axonogenesis. Some key genes in these categories include SRC, VEGFA, EPHA4, DLG4, MAPK3, BMP4, and several semaphorins. These genes can have very different effects on glioblastoma development, with some factors activating oncogenic programs and others behaving as tumor suppressors. Similarly, type I interferon’s effects on treatment are varied. For instance, interferon inhibited proliferation of glioma stem cells and their sphere-forming capacity and induced STAT3 activation [49]. On the other hand, chronic activation of type I IFN signaling has been linked to adaptive resistance to therapy in many tumor types [50].

Activation of oncogenic signals post-radiation could counteract treatment effects and later contribute to relapse. We searched the set of highly up-regulated genes post-radiation for previously identified radio-resistance genes in glioblastoma, oncogenic factors and genes whose high expression is associated with poor prognosis (Table S5). In Table 1, we list these genes according to their molecular function. Since several of these genes have never been characterized in the context of glioblastoma, our results open new opportunities to prevent radio-resistance and increase treatment efficiency. Importantly, there are inhibitors available against several of these proteins (Table S4).

**Table 1.** List of oncogenic factors, genes whose high expression is associated with poor survival and genes previously associated with radio-resistance in GBM that showed increased expression upon radiation. Genes are listed according to molecular function.

| Function | Genes |
| --- | --- |
| Membrane protein | AQP1, ARHGEF2, BAALC, CSF1R, CSPG4, EPS8L2, ERBB3, FGFR4, FYN, GPM6A, ITGB3, JUP |
| Protein kinase | ANKK1, CDKN1A, CSF1R, ERBB3, FAM20C, FGFR4, FYN, IKBKE, MERTK, PDGFRB, SRC, TEC |
| Gene expression regulation | ARID3A, ASAH1, BCL3, BCL6, CBX7, CEBPB, ELF3, FAM20C, FEZF1, HOXA1, HOXB9, JUP, KDM5B, LMO1, LMO2, MACC1, MAF, MSI1, MUC1, NKX2-1, PML, PRDM6, RORC, SATB1, SREBF1, TP53BP1, ZMYM2 |
| Enzymatic activity | ACSS2, AGAP2, APIP, ARHGEF2, C1R, CARD16, CD24, CDKN1A, CEBPB, CSF1R, CSPG4, CTSZ, CUL7, CYTH4, EPS8L2, ERBB3, FAM20C, FGFR4, FTH1, FUCA1, FYN, GHDC, IDO1, IKBKE, ITGB3, JUP, KDM5B, MCF2, MERTK, MFNG, MRAS, NKX2-1,PDE6G,PDGFRB, PML, QPRT, RRM2B, SERPINA5, SFN, SGSH, SRC, SREBF1, TEC, TGFB1, ZMYM2 |
| Phosphotransferase | CSF1R, ERBB3, FAM20C, FYN, IKBKE, MERTK, PDGFRB, SREBF1, TEC |
| Cell surface receptor | BMP7, CSF1R, ERBB3, FGFR4, FYN, ITGB3, ITGB5, LRIG2, MCF2, MERTK, MFNG, PDGFRB, PRDM6, SRC, TEC, TRPM8 |
| Metabolism regulation | ACSS2, APIP, BCL3, BCL6, BTG2, CEBPB,CRTC1, CSPG4, CTSZ, ELF3, FUCA1, IDO1, ITGB3, JUP, MAF, MFNG, NKX2-1, PARP3, PRDM6, PTGES, QPRT, RRM2B, SGSH, TGFB1, TP53BP1, TRPM8, USP9X, VEGFA, ZMYM2 |

### Changes in lncRNA profile in response to radiation

lncRNAs have been implicated in the progression of glioblastoma [51], but their role in response to ionizing radiation is still poorly understood. We identified 161 lncRNAs with expression alterations in T1 vs. T24 comparisons. Analysis of this set with LnC2Cancer [42] identifieddentified several lncRNAs aberrantly expressed in cancer and with relevance to prognosis (Table S1). We also detected significant downregulation of MIR155HG, whose high expression is associated with glioma progression and poor survival [52]. Another downregulated lncRNA with relevance to prognosis is linc000152, whose increased expression has been observed in multiple tumor types [53, 53]. On the other hand, we observed a significant upregulation of two “oncogenic” lncRNAs, NEAT1 and FTX. NEAT1 is associated with tumor growth, grade, and recurrence rate in gliomas [54], while FTX promotes cell proliferation and invasion through negatively regulating miR-342-3p [55]. Thus, if further studies corroborate NEAT1 and FTX as players in radio-resistance, targeting these lncRNAs should be considered to improve treatment response.

### Effect of radiation on splicing

Alternative splicing impacts genes implicated in all hallmarks of cancer [56] and is an important component of changes in expression triggered by ionizing radiation [57]. All types of splicing events (exon skipping, alternative donor, and acceptor splice sites, multiple exclusive exons, and intron retention) were affected similarly upon exposure to radiation (Table S7). At T24, we observed that transcripts associated with RNA-related functions (especially translation), showed the most splicing alterations. Affected transcripts encode ribosomal proteins, translation initiation factors, regulators of translation, and genes involved in tRNA processing and endoplasmic reticulum. Other enriched GO terms include mRNA and ncRNA processing, mRNA degradation, and modification. Catabolism is another process associated with several enriched terms, suggesting that splicing alterations in genes involved in catabolic routes could ultimately contribute to apoptosis (Figure 4A-B, and Table S7). Changes in the splicing profile are likely driven by an alteration in the expression of splicing regulators. In Table 2, we show a list of splicing factors displaying strong expression alterations. Among those previously connected to glioblastoma development upon radiation, LGALS3 is the most extensively characterized. LGALS3 is a galactosidase-binding lectin and non-classic RNA binding protein implicated in pre-mRNA splicing and regulation of proliferation, adhesion, and apoptosis; LGALS3 also is a marker of the early stage of glioma [58].

**Figure 4.**
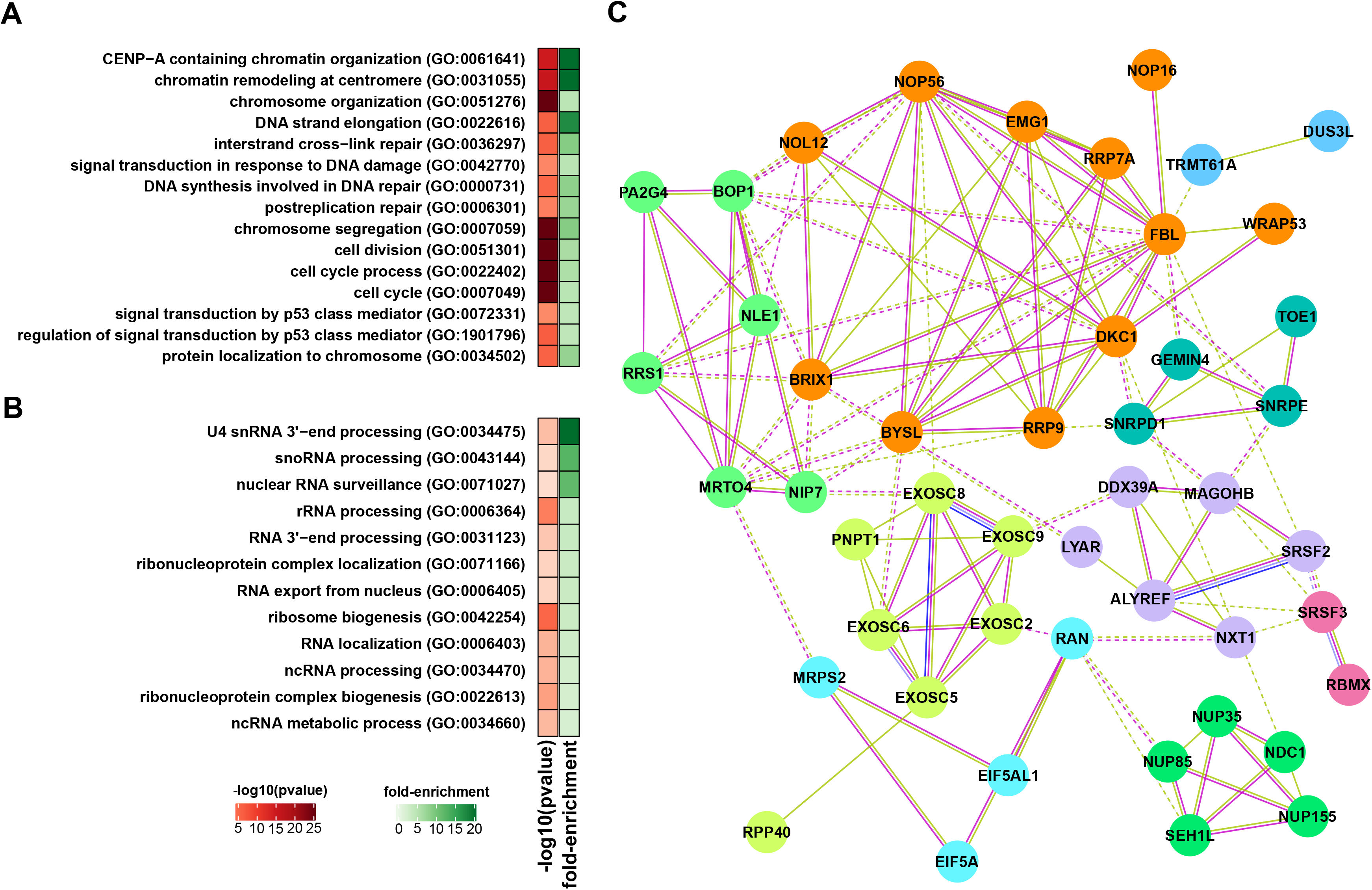
Impact of radiation on the splicing profile of glioblastoma cells. A) GO-enriched terms among genes showing changes in splicing profiles at T24. GO-enriched terms are summarized using REVIGO [38]. B) Protein-protein interaction networks according to STRING [39] showing genes associated with RNA-related functions whose splicing profiles displayed alterations at T24. Gene clusters based on the strength of connection and gene function are identified by color. Lines color indicate type of association: light green, an association based on literature findings; blue indicates gene co-occurrence; magenta indicates experimental evidence.

**Table 2.** Splicing regulators showing changes in expression 24 hours post-radiation. Factors showing an increase in the expression are shown in red, while factors showing a decrease in the expression are represented in blue.

| Gene ID | Gene name | Function |
| --- | --- | --- |
| AHNAK2 | Protein AHNAK2 | splicing regulation |
| ESRP1 | Epithelial splicing regulatory protein 1 | regulation of mRNA splicing |
| LGALS3 | Galectin-3 | signaling receptor binding |
| NOVA2 | RNA-binding protein Nova-2 | alternative splicing regulation |
| SNRPN | Small nuclear ribonucleoprotein N | spliceosomal snRNP assembly |
| ALYREF | THO complex subunit 4 | RNA binding |
| DDX39A | ATP-dependent RNA helicase DDX39A | RNA helicase |
| GEMIN4 | Gem-associated protein 4 | rRNA processing |
| HNRNPL | Heterogeneous nuclear ribonucleoprotein L | alternative splicing regulation |
| LSM2 | U6 snRNA-associated Sm-like protein LSM2 | U6 snRNA-associated Sm-like protein |
| MAGOHB | Mago nashi homolog 2 | exon-exon junction complex |
| PPIH | Peptidyl-prolyl cis-trans isomerase H | ribonucleoprotein complex binding |
| RBMX | RNA-binding motif protein, X chromosome | regulation of mRNA splicing |
| SNRPD1 | Small nuclear ribonucleoprotein Sm D1 | spliceosomal snRNP assembly |
| SNRPE | Small nuclear ribonucleoprotein E | spliceosomal snRNP assembly |
| SRSF2 | Serine/arginine-rich splicing factor 2 | regulation of mRNA splicing |
| SRSF3 | Serine/arginine-rich splicing factor 3 | regulation of mRNA splicing |
| TTF2 | Transcription termination factor 2 | transcription regulation |

### Differential translational efficiency

We used Ribo-seq [59] to identify changes in translation efficiency triggered by radiation. Translation, protein localization, and metabolism appear as top enriched terms among downregulated genes in T1 vs. T24 comparisons (Tables S8-S9). In particular, several ribosomal proteins, along with translation initiation factors and mTOR, showed a significant decrease in translation efficiency (Figure 5A-B). Overall, these results indicate repression of the translation machinery post-radiation exposure and its strong auto-regulation. Since changes in components of the translation machinery are occurring at all levels (transcription, splicing, and translation) at T24, we expect that major translational alterations take place in later stages of post-radiation.

**Figure 5.**
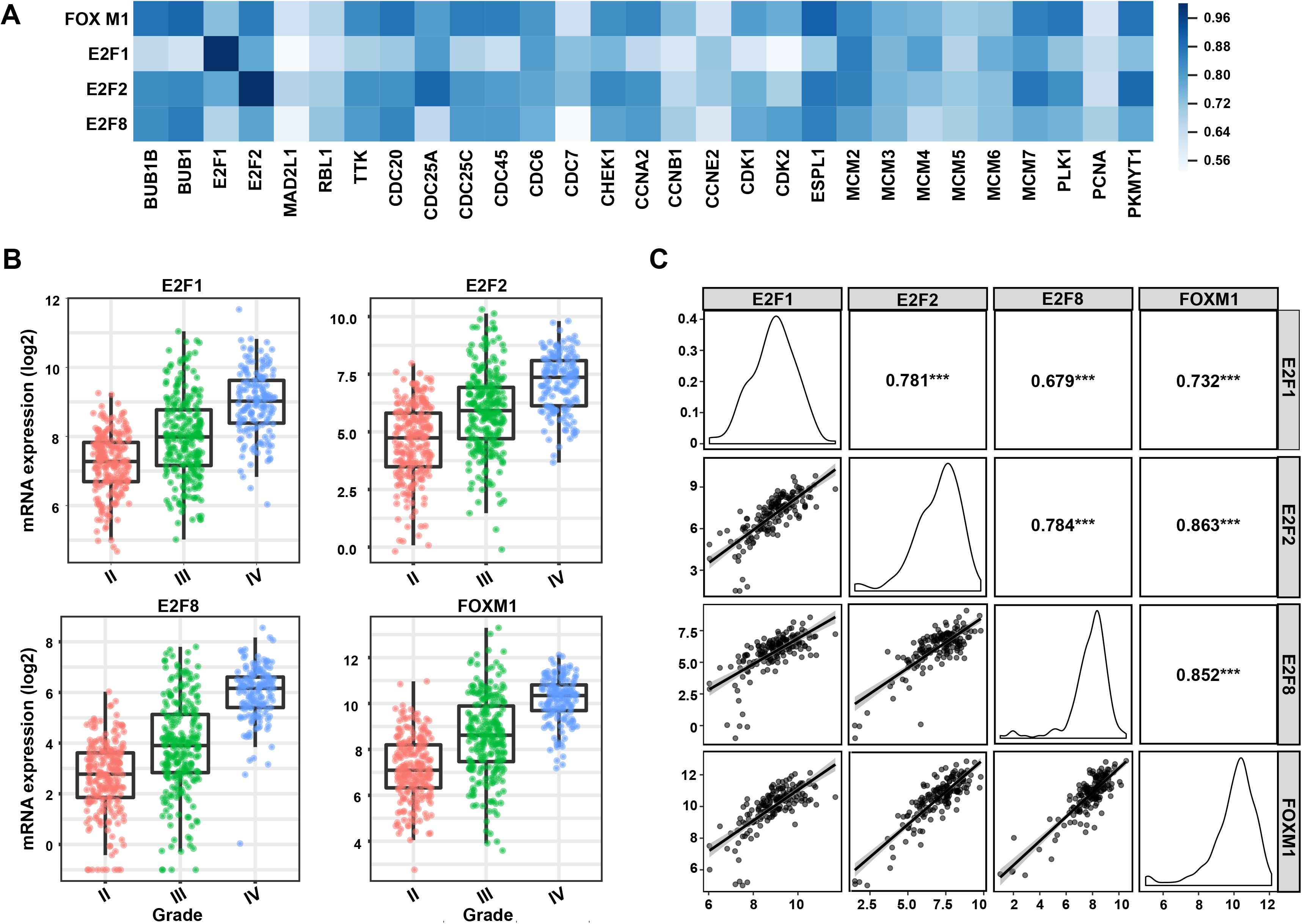
Impact of radiation on the translation profile of glioblastoma cells. A) GO-enriched terms among genes showing changes in translation efficiency at T24. GO-enriched terms are summarized using REVIGO [38]. B) Protein-protein interaction network, according to STRING [39] showing genes whose translation efficiency decreased at T24. Gene clusters based on the strength of connection and gene function are identified by color. Line colors indicate the type of association: light green, an association based on literature findings; blue indicates gene co-occurrence; magenta indicates experimental evidence.

In the upregulated set, we highlight three genes FTH1, APIP, and LRIG2 that could potentially counteract the impact of radiation (Table S10). FTH1 encodes the heavy subunit of ferritin, an essential component of iron homeostasis [60]. Pang *et al*., 2016 [61] reported that H-ferritin plays an important role in radio-resistance in glioblastoma by reducing oxidative stress and activating DNA repair mechanisms. The depletion of ferritin causes down-regulation of ATM, leading to increased DNA sensitivity towards radiation. APIP is involved in the methionine salvage pathway and has a key role in various cell death processes. It can inhibit mitochondria-mediated apoptosis by directly binding to APAF-1 [62]. LRIG2 is a member of the leucine-rich and immunoglobulin-like domain family [63], and its expression levels are positively correlated with the glioma grade and poor survival. LRIG2 promotes proliferation and inhibits apoptosis of glioblastoma cells through activation of EGFR and PI3K/Akt pathway [64].

### Crosstalk between regulatory processes

Parallel analyses of transcription, splicing, and translation alterations in the early response to radiation provided an opportunity to identify crosstalk between different regulatory processes. The datasets showed little overlap, with just a few genes showing alterations in two different regulatory processes. However, we identified several shared GO terms when comparing the results of alternative splicing, mRNA levels, and translation efficiency (Table S10). These terms show two main groups of biological processes. The first group indicates that the expression of genes involved in DNA and RNA synthesis and metabolism is particularly compromised. The second group is related to translation initiation. Ribosomal proteins were particularly affected (Figures 4 and 5). There is growing support for the concept of specialized ribosomes. According to this model, variations in the composition of the ribosome due to the presence or absence of certain ribosomal proteins or alternative isoforms could ultimately dictate which mRNAs get preferentially translated [65]. Therefore, these alterations could later lead to translation changes of a specific set of genes.

## Discussion

We performed the first integrated analysis to define global changes associated with the early response to radiation in glioblastoma. Our approach allowed the identification of “conserved” alterations at the transcription, splicing and translation levels and defined possible crosstalk between different regulatory processes. Alterations at the level of transcription were dominant, but changes affecting genes implicated in RNA mediated regulation were ubiquitous; they indicate that these processes are important components in radio-response and suggest that more robust changes in splicing and translation might take place later.

### E2F1, E2F2, E2F8, and FOXM1 as major drivers of transcriptional responses upon radiation

We observed marked changes in the mRNA levels of genes implicated in cell cycle, DNA replication, and repair 24 hours (T24) after radiation. Downregulation of several transcription factors, most of them members of the zinc finger family, was observed at one hour post-radiation. This group displays high correlations in expression within glioblastoma samples from TCGA, suggesting that they might work together to regulate gene expression. Unfortunately, most are poorly characterized, and the lack of information has prevented establishing further connections to changes in the cell cycle and DNA replication that we observed at T24.

GSEA and expression correlation analysis suggested that the downregulation of members of the E2F family is likely responsible for several of the expression changes we observed at T24. E2Fs have been defined as major transcriptional regulators of the cell cycle. The family has eight members that could act as activators or repressors depending on the context, and are known to regulate one another. They are upregulated in many tumors due to overexpression of cyclin-dependent kinases (CDKs), inactivation of CDK inhibitors, or RB Transcriptional Corepressor 1 (RB1) and are linked to poor prognosis. Alterations in E2F genes can induce cancer in mice [66, 67]. Specifically, we found that three E2F members showed decreased expression upon radiation: E2F1, E2F2, and E2F8, all of which have been previously implicated in glioblastoma development.

E2F1 is probably the best-characterized member of the E2F family. Besides its known effect on cell cycle regulation and DNA replication, it is also a positive regulator of telomerase activity, binding the TERT promoter [68]. Recent studies show that lncRNAs and miRNAs function in an antagonistic fashion to regulate E2F1 expression, ultimately affecting cell proliferation, glioblastoma growth, and response to therapy [69, 70, 71]. E2F2 has been linked to the maintenance of glioma stem cell phenotypes and cell transformation [72, 73]. Several tumor suppressor miRNAs (let7b, miR-125b, miR-218, and miR-138) decrease the proliferation and growth of glioblastoma cells by targeting E2F2 [72, 74, 75, 76]. Although still poorly characterized in the context of glioblastoma, E2F8 drives an oncogenic phenotype in glioblastoma. Its expression is modulated by HOXD-AS1, which serves as a sponge and prevents the binding of miR-130a to E2F8 transcripts [77]. FOXM1 is another potential regulator of the group of cell cycle and DNA replication genes affected by radiation. FOXM1 is established as an important player in chemo- and radio-resistance and a contributor to glioma stem cell phenotypes [5, 6, 78, 79, 80, 81, 82, 83]. FOXM1 and E2F protein have a close relationship and share target genes [84]. Additionally, FOXM1- and E2F2-mediated cell cycle transitions are implicated in the malignant progression of IDH1 mutant glioma [85].

E2F and FOXM1 targeting could be considered as an option to increase radio-sensitivity. Since the development of transcription factor inhibitors is very challenging, an alternative to be considered is the use of BET (bromodomain and external) inhibitors. BET is a family of proteins that function as readers for histone acetylation and modulates the transcription of oncogenic programs [86]. Recent studies in glioblastoma with a new BET inhibitor, dBET6, showed promising results and established that its effect on cancer phenotypes comes via disruption of the transcriptional program regulated by E2F1 [87].

### RNA processing and regulation as novel categories in radio-response

Besides the expected changes in expression of cell cycle, DNA replication and repair genes, radiation affected preferentially the expression of genes implicated in RNA processing and regulation. Additionally, we identified a co-expression module containing multiple genes associated with translation initiation, rRNA and snoRNA processing, RNA localization, and ribonucleoprotein complex biogenesis.

Many regulators of RNA processing are implicated in glioblastoma development, and splicing alterations affect all hallmarks of cancer [88]. Radiation-induced changes in the splicing patterns of oncogenic factors and tumor suppressors such as CDH11, CHN1, CIC, EIF4A2, FGFR1, HNRNPA2B1, MDM2, NCOA1, NUMA1, RPL22, SRSF3, TPM3, APC, CBLB, FAS, PTCH1, and SETD2. We also observed changes in expression of four RNA processing regulators previously identified in genomic/functional screening for RNA binding proteins contributing to glioblastoma phenotypes: MAGOH, PPIH, ALYREF, and SNRPE [89].

### Potential new targets to increase radio-sensitivity and prevent relapse

Activation of oncogenic signals is an undesirable effect of radiation that could influence treatment response and contribute to relapse. We observed increased expression or translation and splicing alterations of a number of pro-oncogenic factors, genes whose high expression is associated with poor survival and genes previously implicated in radio-resistance.

Among genes with the most marked increase in expression upon radiation, we identified members of the Notch pathway (HES2, NOTCH3, MFNG, and JAG2). Notch activation has been linked to radio-resistance in glioblastoma, and Notch targeting improves the results of radiation treatment [90, 91]. We also identified several genes associated with the PI3K-Akt, Ras, and Rap1 signaling pathways that increased expression levels upon radiation exposure. Targeting these pathways has been explored as a therapeutic option in glioblastoma [92, 90]. Other oncogenic factors relevant to glioblastoma that had increased expression after radiation exposure include SRC, MUC1, LMO2, PML, PDGFR*β*, BCL3, and BCL6.

Anti-apoptotic genes (BCL6, RRM2B, and IDO1) also showed increased expression upon radiation. BCL6 is a member of the ZBTB family of transcription factors, which functions as a p53 pathway repressor. The blockage of the interaction between BCL6 and its cofactors has been established as a novel therapeutic route to treat glioblastoma [93]. RRM2B is an enzyme essential for DNA synthesis and participates in DNA repair, cell cycle arrest, and mitochondrial homeostasis. The depletion of RRM2B resulted in ADR-induced apoptosis, growth inhibition, and enhanced sensitivity to chemo- and radiotherapy [94]. IDO1 is a rate-limiting metabolic enzyme involved in tryptophan metabolism that is highly expressed in numerous tumor types [95]. The combination of radiation therapy and IDO1 inhibition enhanced therapeutic response [96].

Among genes whose high expression correlates with decreased survival in glioblastoma, we identified several components of the “matrisome” and associated factors (FAM20C, SEMA3F, ADAMTSL4, ADAMTS1 SERPINA5, and CRELD1). The core of the “matrisome” contains ECM proteins, while associated proteins include ECM-modifying enzymes and ECM-binding growth factors. This complex of proteins assembles and modifies extracellular matrices, contributing to cell survival, proliferation, differentiation, morphology, and migration [97]. In addition, several genes of the proteinase inhibitor SERPIN family (SERPINA3, SERPINA12, SERPINA5, and SERPINI1) implicated in ECM regulation [98] were among those with high levels of expression upon radiation.

## Conclusions

In conclusion, our results generated a list of candidates for combination therapy. Contracting the effect of oncogenic factors and genes linked to poor survival could increase radio-sensitivity and treatment efficiency. Importantly, there are known inhibitors against several of these proteins (Table S5). Moreover, RNA processing and translation were determined to be important components of radio-response. These additional vulnerable points could be explored in therapy, as many inhibitors against components of the RNA processing and translation machinery have been identified [99, 100].

## List of abbreviations

TCGA: The Cancer Genome Atlas
NSCs: Neural stem cells
lncRNAs: long non-coding RNAs
Ribo-seq: high-throughput ribosome profiling
T0: time point corresponding to no irradiation
T1: time point corresponding to 1 hour post irradiation
T24: time point corresponding to 24 hours post irradiation
CDS: coding domain sequence
PCA: Principle component analysis
BH: Benjamini and Hochberg FDR adjustment procedure
WGCNA: Weighted Gene co-expression network analysis
TPM: Transcripts per million
kME: eigene-gene based connectivity in cluster analysis
GO: Gene ontology
GSEA: Gene set enrichment analysis

## Ethics approval and consent to participate

Not applicable.

## Consent for publication

All authors give their consent for publication.

## Availability of data and materials

The processed data of read abundance matrices is available through GEO accession GSE141013. Scripts for differential expression analysis and translational efficiency analysis are available at https://github.com/saketkc/2019_radiation_gbm

## Author’s contributions

SC. Contributed to experimental design, conducted most of the data analysis, manuscript writing.

SCB. Performed all experiments.

HM. Contributed to data analysis, manuscript writing.

WL. Contributed to data analysis.

DV. Contributed to data interpretation and writing.

MQ. Contributed to the experimental part.

ADS. Design of analysis pipeline.

LOFP. Experimental design and concept, data analysis, manuscript writing.

## Additional Files

Additional file 1: Figure S1. Experimental design and Principal Component Analysis of RNA-Seq and Ribo-Seq data from glioblastoma cell lines.

A) Schematic representation of experimental protocol followed for radiation exposure of glioma cell lines, and sample preparation for sequencing the RNA and ribosome footprints. B) Principal component analyses performed on normalized log-transformed read counts of RNA-Seq and Ribo-Seq datasets.

Additional file 2: Figure S2. Fragment length distribution of ribosome footprints of glioblastoma cell lines.

Fragment lengths distribution was obtained using ribotricer.

Additional file 3: Figure S3. The ribosome density profiles of glioblastoma cell lines.

The metagene distributio was obtained using ribotricer.

Additional file 4: Figure S4. Global view of glioblastoma cell lines transcription and translation profiles after radiation

A) Differentially expressed genes (left) and the number of genes whose translation efficiency is differentially regulated (right) after radiation exposure at different time points. B) Volcano plots showing the expression and transition alterations of genes at 24 hr compared to 1 hr after radiation exposure. Blue dots indicate upregulated genes (adjusted p-value ¡ 0.05, log2 fold change < 0), and orange dots indicate downregulated genes. (adjusted p-value < 0.05, log2 fold change < 0). C) Sizes of gene modules found in U251 and U343 cell lines. D) Preservation Median Rank and *Z*_summary_ for all modules. A lower median rank indicates the module is preserved, and the corresponding modules in U251 and U343 cell lines share a high number of genes. A *Z*_summary_ score of 2-10 indicates weak preservation, while a *Z*_summary_ > 10 indicates high preservation.

Additional file 5: Table S1. Differentially expressed genes at different time points after radiation exposure in two glioblastoma cell lines.

Genes were tested for differential expression across two conditions using DESeq2 (v1.16.1). (Sheet 1) Differentially expressed protein-coding genes at baseline versus 1 hour after radiation. (sheet 2) Transcription factors showing high expression correlation in TCGA glioblastoma samples. Data were analyzed with Gliovis. (Sheet 3) Differentially expressed protein-coding genes at T24vsT1. (Sheet 4) Differentially expressed lncRNAs at T1vsT24. (Sheet 5) lincRNAs known to be connected to cancer.

Additional file 6: Table S2. Functional enrichment analysis for genes with differential expression at different time points after radiation exposure in two glioblastoma cell lines.

Functional enrichment and pathway analyses were performed using Panther and Reactome databases, respectively. (Sheet 1) Gene Ontology analysis of downregulated genes at 1 hr versus 24 hr after radiation exposure. (Sheet 2) Pathway analysis of downregulated genes at 1 hr versus 24 hr after radiation exposure. (Sheet 3) Gene Ontology analysis of upregulated genes at 1 hr versus 24 hr after radiation exposure. (Sheet 4) Pathway analysis of upregulated genes at 1 hr versus 24 hr after radiation exposure.

Additional file 7: Table S3. List of gene modules in U251 and U343 cell lines and their corresponding genes.

WGCNA was used for identifying tightly regulated gene modules. (Sheet 1) U251. (Sheet 2) U343 (Sheet 3) List of protein-coding hub genes in gene modules 2. (Sheet 3) Enriched GO terms in module 2. (Sheet 4) Enriched Pathways in Module 2.

Additional file 8: Table S4. FOXM1, E2F1, E2F2, and E2F8 are potential regulators of genes showing a decrease in expression upon radiation. (Sheets 1-4) Genes showing high expression correlation with FOXM1, E2F1, E2F2, and E2F8 in TCGA glioblastoma samples, according to Gliovis. (Sheet 5) Downregulated genes displaying high expression correlation with FOXM1, E2F1, E2F2 and E2F8 in TCGA glioblastoma samples. (Sheet 6) Gene ontology analysis of the gene set in Sheet 5.

Additional file 9: Table S5. Oncogenes, genes contributing to radio-resistance and genes linked to GBM survival that are upregulatd upon radiation exposure.

(Sheets 1-4) List of inhibitors against relevant genes. (Sheet 5) Oncogenes, genes contributing to radio-resistance and genes linked to GBM survival that are upregulatd upon radiation exposure.

Additional file 10: Table S6. List of splicing events and their respective genes affected by radiation.

(Sheets 1 and 2) Splicing analysis was performed using rMATS (v4.0.1) across T24, T1, and T0 time points. Splicing events are as defined in rMATS. (Sheet 3) RNA processing and translation genes showing changes in splicing post-radiation.

Additional file 11: Table S7. Functional enrichment analysis for genes displaying changes in splicing profile upon radiation.

Functional enrichment and Pathway analysis were performed using the Panther and Reactome databases respectively. (Sheet 1) Gene Ontology. (Sheet 2) Pathway analysis.

Additional file 12: Table S8. Genes displaying changes in translation efficiency after radiation exposure.

Translationally efficient genes discovered using Riborex (v1.2.3).

Additional file 13: Table S9. Functional enrichment analysis for genes with differential translational efficiency after radiation exposure.

Functional enrichment and Pathway analysis were performed using the Panther and Reactome databases, respectively. (Sheet 1) Gene Ontology. (Sheet 2) Pathway analysis.

Additional file 14: Table S10. Comparison of GO terms in the different analyses.

(Sheet 1) Analysis shows enriched GO terms common to genes showing changes in splicing profile, with decreased translational efficiency, and decreased mRNA levels after radiation exposure. Only shared GO terms are shown. Colors reflect the studies sharing the terms. (Sheet 2) Two main categories of biological processes shared by the three different analyses.

