## Supplementary figures and images for "Integrated analyses of early responses to radiation in glioblastoma identify new alterations in RNA processing and candidate target genes to improve treatment outcomes"

### Figure S1

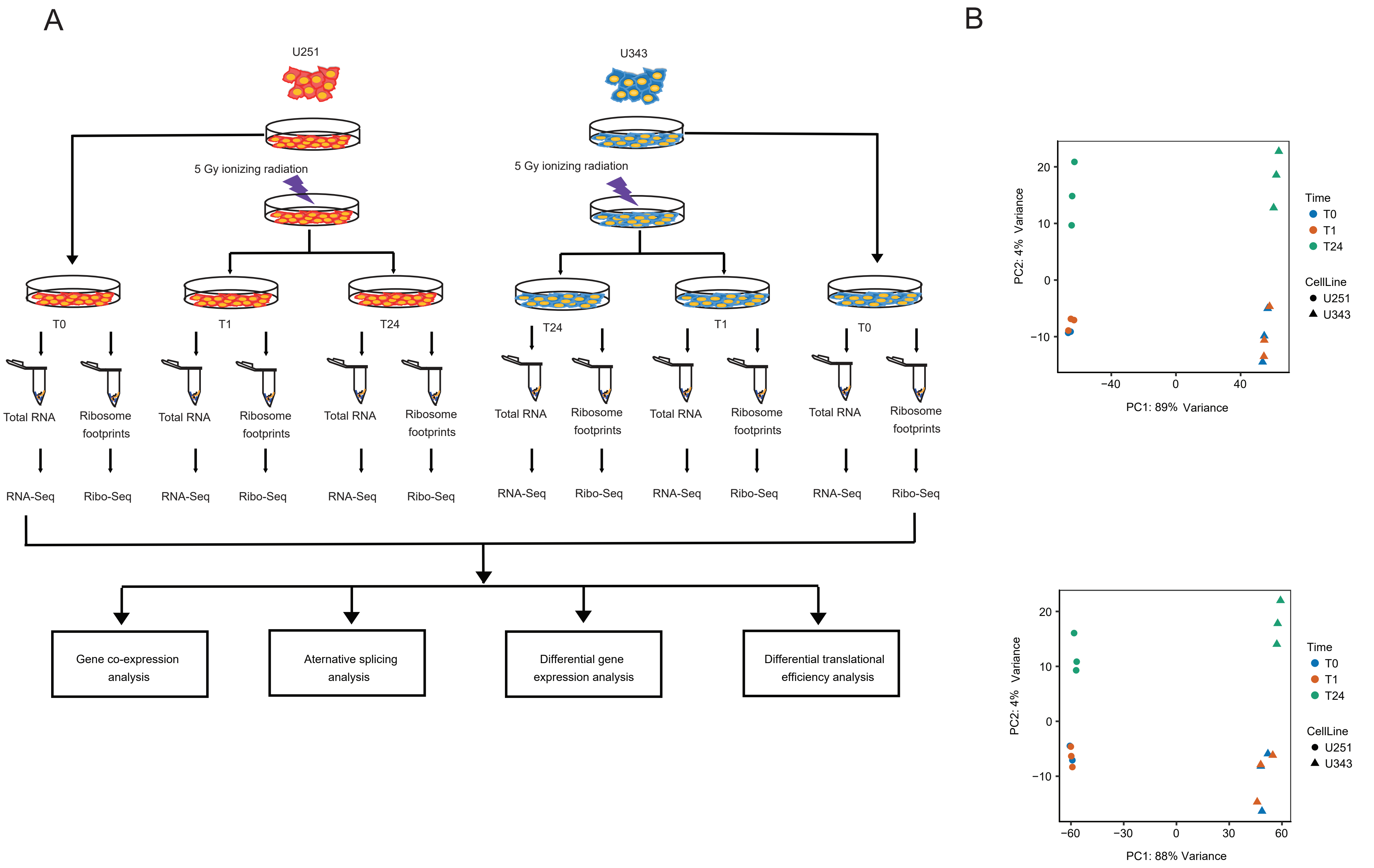

### Figure S2

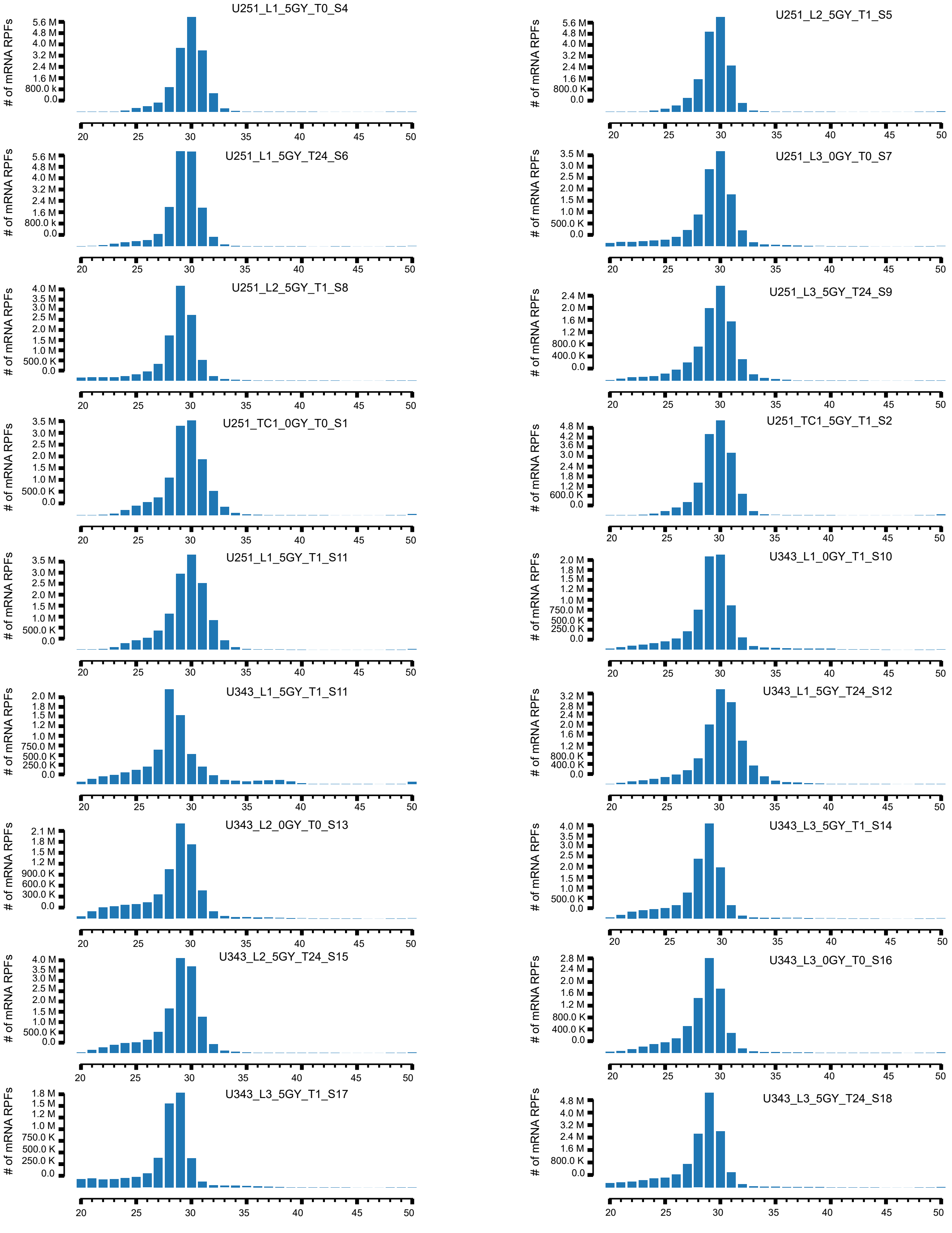

### Figure S3

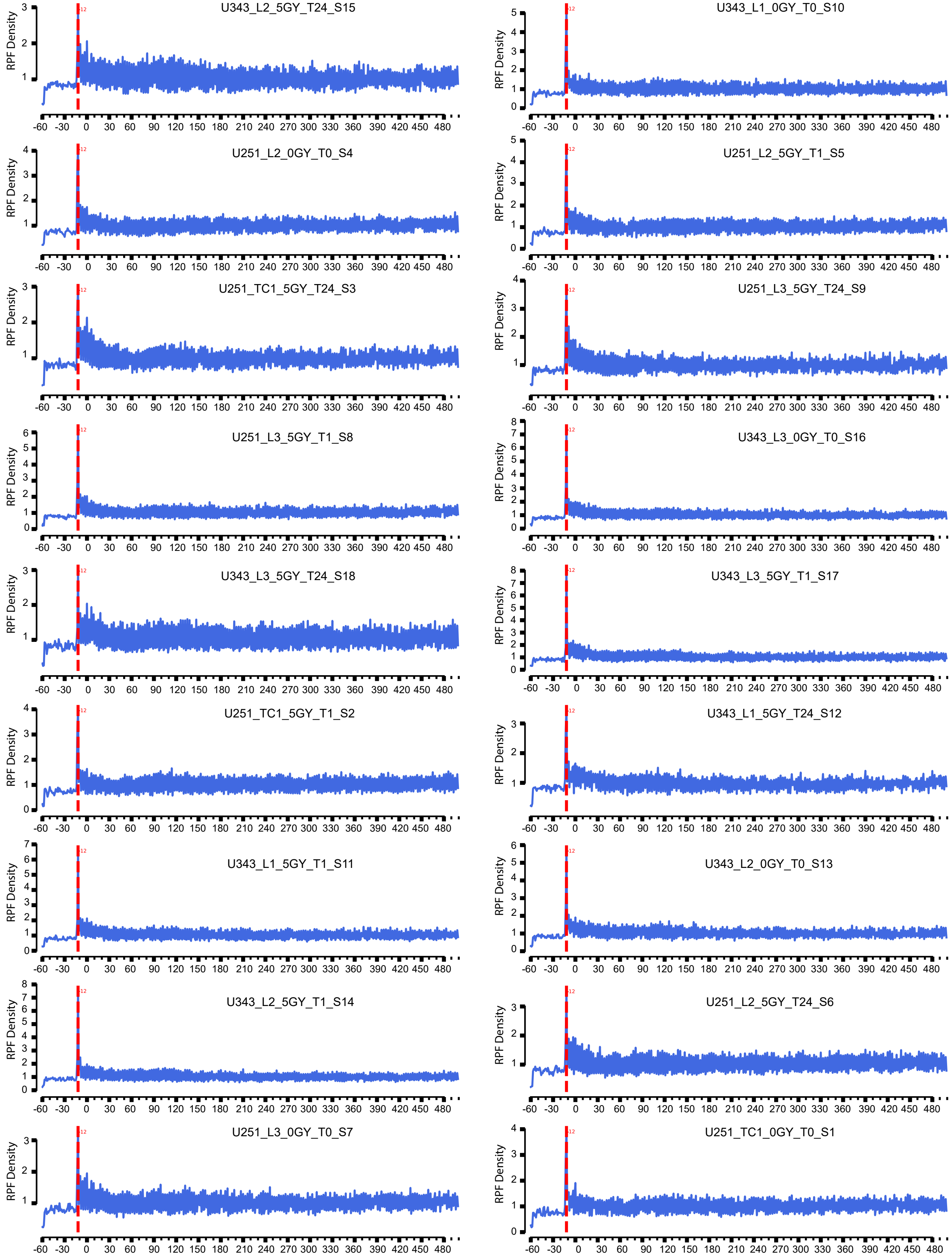

### Figure S4

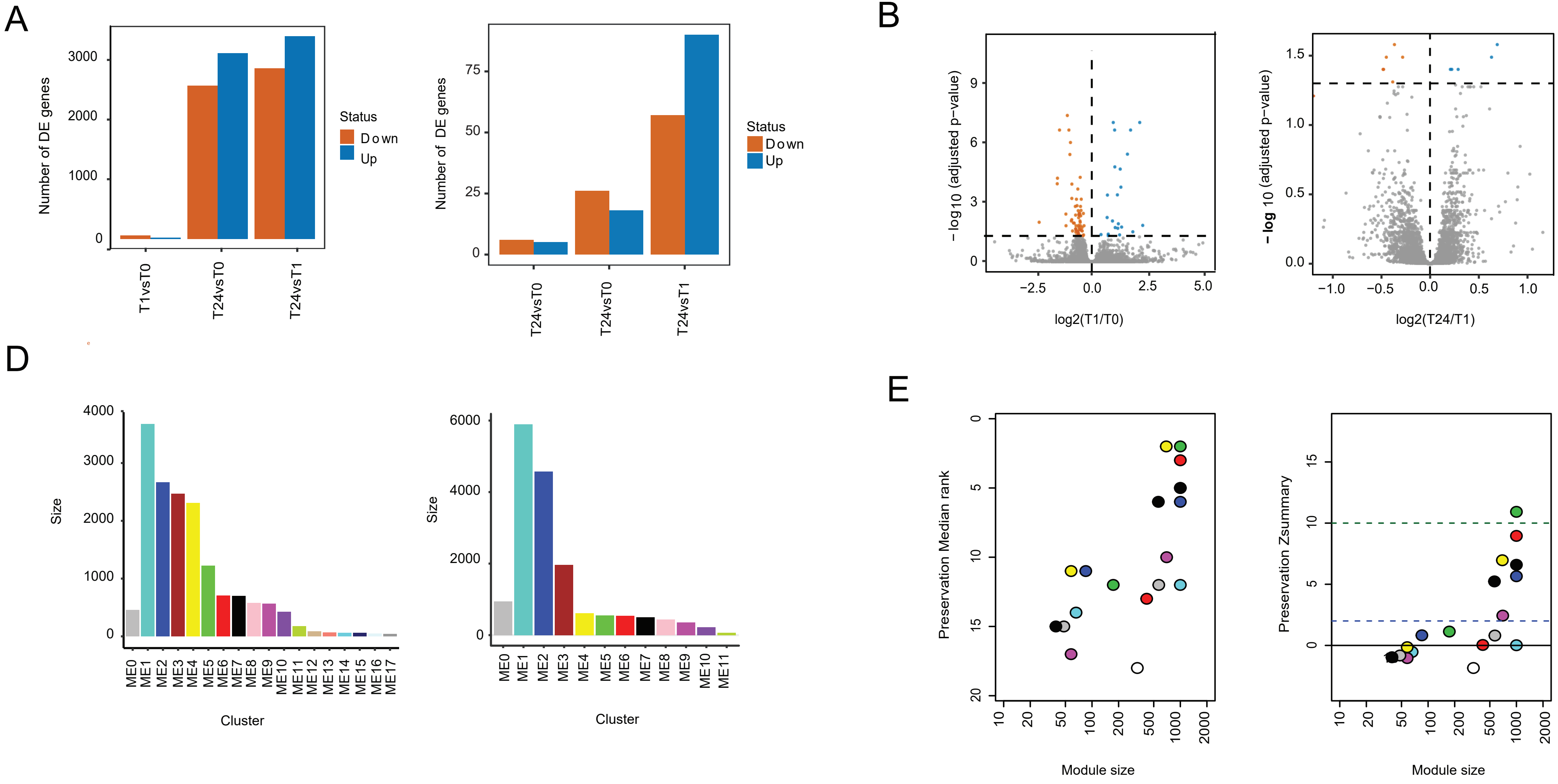
