## Supplemental Data 1 for "Integrated analyses of early responses to radiation in glioblastoma identify new alterations in RNA processing and candidate target genes to improve treatment outcomes"

**Figure S1. Experimental design and Principal component analysis of RNA-Seq and Ribo-Seq data from glioblastoma cell lines. A)** Schematic representation of experimental protocol followed for radiation of glioma cell lines, and sample preparation for sequencing the RNA and ribosome footprints. **B)** Principal component analyses performed on normalized log‐transformed read counts of RNA-Seq and Ribo-Seq datasets.

**Figure S2. Fragment length distribution of ribosome footprints from glioblastoma cell lines.**

**Figure S3. The ribosome density profiles from glioblastoma cell lines.**

**Figure S4. Global view of glioblastoma cell lines transcription and translation profiles after radiation. A)** Barplots showing the number of differentially expressed genes (left) and the number of genes whose translation efficiency is differentially regulated (right), upon radiation at different time points. **B)**: Volcano plots showing the expression and transition alterations of genes in T24 compared to T1 upon radiation. The blue dots indicate upregulated genes (Adjusted p-value < 0.05, log_2_ fold change > 0), and orange dots indicate downregulated genes. (Adjusted p-value < 0.05, log_2_ fold change < 0). **C)** Size of gene modules found in U251 and U343 **D)** Preservation Median Rank and Zsummary for all modules. A lower median rank indicates the module is preserved and the corresponding modules in U251 and U343 share high number of genes. A Zsummary score of 2-10 indicates weak preservation while a Zscore>10 indicates high preservation.
